# Combining Blood-Based Biomarkers and Structural MRI Measurements to Distinguish Persons With and Without Significant Amyloid Plaques

**DOI:** 10.1101/2023.10.20.563332

**Authors:** Yanxi Chen, Yi Su, Jianfeng Wu, Kewei Chen, Alireza Atri, Richard J Caselli, Eric M. Reiman, Yalin Wang, the Alzheimer’s Disease Neuroimaging Initiative

**Author notes:** **Please address correspondence to:** Dr. Yalin Wang, School of Computing and Augmented Intelligence, Arizona State University, P.O. Box 878809, Tempe, AZ 85287 USA, **E-mail:**. Acknowledgments: Data used in preparing this article were obtained from the Alzheimer’s Disease Neuroimaging Initiative (ADNI) database (adni.loni.usc.edu). As such, many investigators within the ADNI contributed to the design and implementation of ADNI and/or provided data but did not participate in the analysis or writing of this report. A complete listing of ADNI investigators can be found at: http://adni.loni.usc.edu/wp-content/uploads/how_to_apply/ADNI_Acknowledgement_List.pdf.

## Abstract

**Background:** Amyloid-β (Aβ) plaques play a pivotal role in Alzheimer’s disease. The current positron emission tomography (PET) is expensive and limited in availability. In contrast, blood-based biomarkers (BBBMs) show potential for characterizing Aβ plaques more affordably. We have previously proposed an MRI-based hippocampal morphometry measure to be an indicator of Aβ-plaques.

**Objective:** To develop and validate an integrated model to predict brain amyloid PET positivity combining MRI feature and plasma Aβ42/40 ratio.

**Methods:** We extracted hippocampal multivariate morphometry statistics (MMS) from MR images and together with plasma Aβ42/40 trained a random forest classifier to perform a binary classification of participant brain amyloid PET positivity. We evaluated the model performance using two distinct cohorts, one from the Alzheimer’s Disease Neuroimaging Initiative (ADNI) and the other from the Banner Alzheimer’s Institute (BAI), including prediction accuracy, precision, recall rate, F1 score and AUC score.

**Results:** Results from ADNI (mean age 72.6, Aβ+ rate 49.5%) and BAI (mean age 66.2, Aβ+ rate 36.9%) datasets revealed the integrated multimodal (IMM) model’s superior performance over unimodal models. The IMM model achieved prediction accuracies of 0.86 in ADNI and 0.92 in BAI, surpassing unimodal models based solely on structural MRI (0.81 and 0.87) or plasma Aβ42/40 (0.73 and 0.81) predictors.

**Conclusion:** Our IMM model, combining MRI and BBBM data, offers a highly accurate approach to predict brain amyloid PET positivity. This innovative multiplex biomarker strategy presents an accessible and cost-effective avenue for advancing Alzheimer’s disease diagnostics, leveraging diverse pathologic features related to Aβ plaques and structural MRI.

## 1 INTRODUCTION

Clinical psychiatrist and neuroanatomist Alois Alzheimer first reported the symptoms of a “pre-senile” and thought to be relatively rare dementia associated with abnormal brain plaque and neurofibrillary emergence and aggregation in November 1906 [1]. However, over that last century, Alzheimer’s disease (AD) has been recognized as a major global health concern that is primarily causing and contributing to cognitive impairment and dementia worldwide with increasing burden and costs [2,3]. While no cure for AD exists, recent therapeutic advances allow for utilization of both symptomatic and disease-modifying treatments across the AD clinical spectrum to slow clinical progression, thus making the timely diagnosis of AD more important than ever [4–9].

Brain imaging modalities currently serve as primary methods for assessment of dementia and AD related brain abnormalities, including for the presence of amyloid and tau pathology [10–12]. Among them, magnetic resonance imaging (MRI) detects brain shrinkage [13], while positron emission tomography (PET) tracers can measure amyloid-plaque and tau neurofibrillary tangle burden, respectively [12,14,15]. Although PET scans have demonstrated high accuracy in detecting AD related pathological changes, their costs, radiation exposure, and insurance coverage status restrict their widespread clinical use and can also limit their use in clinical trials [8,12]. As a less invasive, less costly, and more readily available modality that is considered part of the standard clinical diagnostic pathway [6,16], MRI has the potential to characterize brain atrophy patterns and changes over time [13]; and can also characterize other useful structural (and functional) features beyond basic volumetric measures.

We have previously developed a series of advanced structural MRI methods and demonstrated their potential advantages in AD diagnostics including multivariate morphometry statistics (MMS) [17] which uses multivariate tensor-based morphometry (mTBM) to encode surface morphometry along the surface tangent direction and radial distance (RD) to encode morphometry along the surface normal direction. This approach outperformed volumetric measures in detecting hippocampal deformation and also exhibited superior performance in determining group differences between AD diagnosis groups [18,19]. To address the high dimensionality problem of this method in classification tasks, we developed patch analysis-based surface sparse-coding and max-pooling (PASS-MP) system to extract patch-based image features and learn a low-dimensional representation of hippocampal MMS that are predictive of longitudinal cognitive decline [20]. Furthermore, the objective function was modified to improve the sparse coding system by integrating the correntropy based method [21].

Some studies have suggested that it may be feasible to impute amyloid positivity from MRI, potentially providing a lower cost and more accessible alternative compared to more accurate but costly PET scans [22]. For example, Duygu et al. used combined structural MRI and cerebral blood flow (CBF) patterns as predictors of amyloid positivity status and achieved 83% classification accuracy in early MCI individuals [22]. Leveraging the advanced MR analysis techniques our team has previously developed [17–21], we also demonstrated the feasibility of potentially predicting amyloid burden based on MR data [23].

Recently, there is an increased interest and success in the development of blood-based biomarkers (BBBMs) as research and diagnostic tools for the characterization of amyloid and tau pathologies [24–26]. In a recent review of using BBBMs in detecting amyloid positivity [27], the highest prediction accuracy using Aβ42/40 as the predictor was 0.77 [28]. Another study of using plasma Aβ42/40 ratio as a biomarker for amyloidosis reported an area under the ROC curve (AUC-ROC) of 0.81 (accuracy 75%). The performance was further improved after controlling for cohort heterogeneity (AUC-ROC 0.86 and accuracy 81%) and adding APOE4 copy number and participant age (AUC-ROC 0.90 and accuracy 86%) indicating a high diagnostic accuracy in detection of brain amyloidosis [29]. Also, a recently proposed method combining MRI with BBBMs for longitudinal structural atrophy and cognitive decline prediction demonstrated reliable performance in distinguishing fast vs slow progressors in MCI [30].

In this study, we propose a novel pipeline that integrates imaging features from MRI with an AD BBBM to predict PET-based brain Aβ positivity. We hypothesized that by combining information from structural MRI and an Aβ BBBM in an integrated multimodal (and multiplex) model prediction of amyloid PET positivity would be improved compared to performance by each unimodal model.

## 2 MATERIALS and METHODS

### 2.1 Data Acquisition and Preprocessing

In this study, we identified and downloaded data in 198 subjects reported in a previous study (Table 1) [31] from the ADNI database with matching plasma Aβ42/40 measures, florbetapir PET, and T1-MRI scans. In addition, we obtained 111 subjects from Banner Alzheimer’s Institute (BAI) with [^11^C]-Pittsburgh compound B (PiB) PET, T1-MRI, and plasma Aβ42/40 measures [29]. Plasma Aβ42/40 ratios in ADNI and BAI datasets were both quantified using a LC-MS based assay [29,31]. AUC scores of 0.89 in the ADNI cohort and 0.81 in the BAI cohort were reported respectively by the previous studies [29,31], indicating promising and consistent performance of this assay to detect brain amyloid positivity. Image analysis was done using previously established methods [32,33]. Briefly, T1-MRI were processed with FreeSurfer, PET images were motion corrected, summed, and co-registered with MRI, regional standard uptake value ratio (SUVR) with and without partial volume correction was calculated using cerebellar cortex as the reference region, and target region for global index of amyloid burden (mean cortical SUVR, MCSUVR) included frontal, parietal, temporal, and precuneus regions [32,33]. The Centiloid procedure was adopted to harmonized PET derived amyloid burden measure between the two studies using the two different tracers [34]. According to this procedure, as we previously established, the equivalent amyloid positivity thresholds adopted were MCSUVR > 1.42 for PiB and MCSUVR > 1.19 for florbetapir, with the positivity threshold translated to approximately 16 Centiloid unit [35,36]. Based on this criteria, 98 subjects were amyloid positive and 100 were amyloid negative for the ADNI cohort, and 41 were Aβ positive and 70 were Aβ negative for the BAI cohort. An overview of the participant summary statistics can be seen in Table 1.

**Table 1.** Summary statistics of the patients.

|  | ADNI (N = 198) | BAI (N = 111) |
| --- | --- | --- |
| **Age** |  |  |
| Min | 55.7 | 47.0 |
| Max | 88.4 | 85.0 |
| Mean ± SD | 72.62 ± 6.92 | 66.21 ± 7.95 |
| **Sex** |  |  |
| Male | 94 (47%) | 27 (24.3%) |
| Female | 103 (52%) | 84 (75.7%) |
| **Diagnosis** |  |  |
| CN | 106 (53.5%) | 111 (100%) |
| MCI | 92 (46.5%) | 0 (0%) |
| **APOE4** |  |  |
| NC | 122 (61.6%) | 55 (49.5%) |
| HT | 68 (34.3%) | 35 (31.5%) |
| HM | 8 (4.04%) | 21 (18.9%) |
| **Aβ positivity** |  |  |
| Aβ+ | 91 (49.5%) | 41 (36.9%) |
| Aβ- | 107 (50.5%) | 70 (63.1%) |

### 2.2 System Overview

Our prediction pipeline can be divided into two parts: 1) Hippocampal structure segmentation and registration: The hippocampal structures were segmented and registered from structural MRI (sMRI). Vertex-wise MMS features were computed for both hemispheres of the brain. These MMS features consist of Radial Distance (RD) and multivariate tensor-based morphometry (mTBM), measuring hippocampal size and the deformation within the surface, respectively; 2) Patch-based feature extraction and classification: Patches were selected from the hippocampal surfaces and sparse codes were generated using the correntropy-induced sparse coding method. Subsequently, a max-pooling step was applied to reduce the dimensionality to a reasonable scale for subsequent machine learning algorithms. The resulting feature vectors were input into a random forest classifier both by themselves and combining with plasma Aβ42/40 features.

By employing this pipeline, we aimed to leverage the unique information provided by the hippocampal MMS features and the discriminative power of the correntropy-induced sparse coding method to enhance the predictive performance. Additionally, we investigated the potential improvement achieved by integrating the image-based features with the plasma Aβ42/40 levels.

### 2.3 Hippocampus Segmentation and Surface Registration

We took advantage of our previously established pipeline protocol for hippocampus segmentation and image registration [14,28,29], which employs FMRIB’s Integrated Registration and Segmentation Tool (FIRST) for automatic segmentation, a topology-preserving level set method combined with marching cubes algorithm for hippocampal surface reconstruction [37–39].[30][31][32]We also performed surface smoothing to reduce the noise, followed by mesh simplification and refinement using the same method as our previous work [40–42]. A conformal grid of size 150*100 was computed on each surface using a holomorphic 1-form basis [43,44], followed by the computation of conformal factor and mean curvature on each point and a surface fluid registration method to register the hippocampal surfaces to a template surface [45–47][40] For more details, refer to [45].

### 2.4 Surface Multivariate Morphometry Statistics

After the surface registration was completed, we computed RD and mTBM as surface features. The RD measures the hippocampal size by the surface differences along the surface normal direction, while mTBM measures the local surface deformation and has a relatively stronger signal detection power [40,48,49], resulting in a feature vector of size 4*1 for each of the 15000 vertices on each hemisphere (Figure 1).

**Figure 1.**
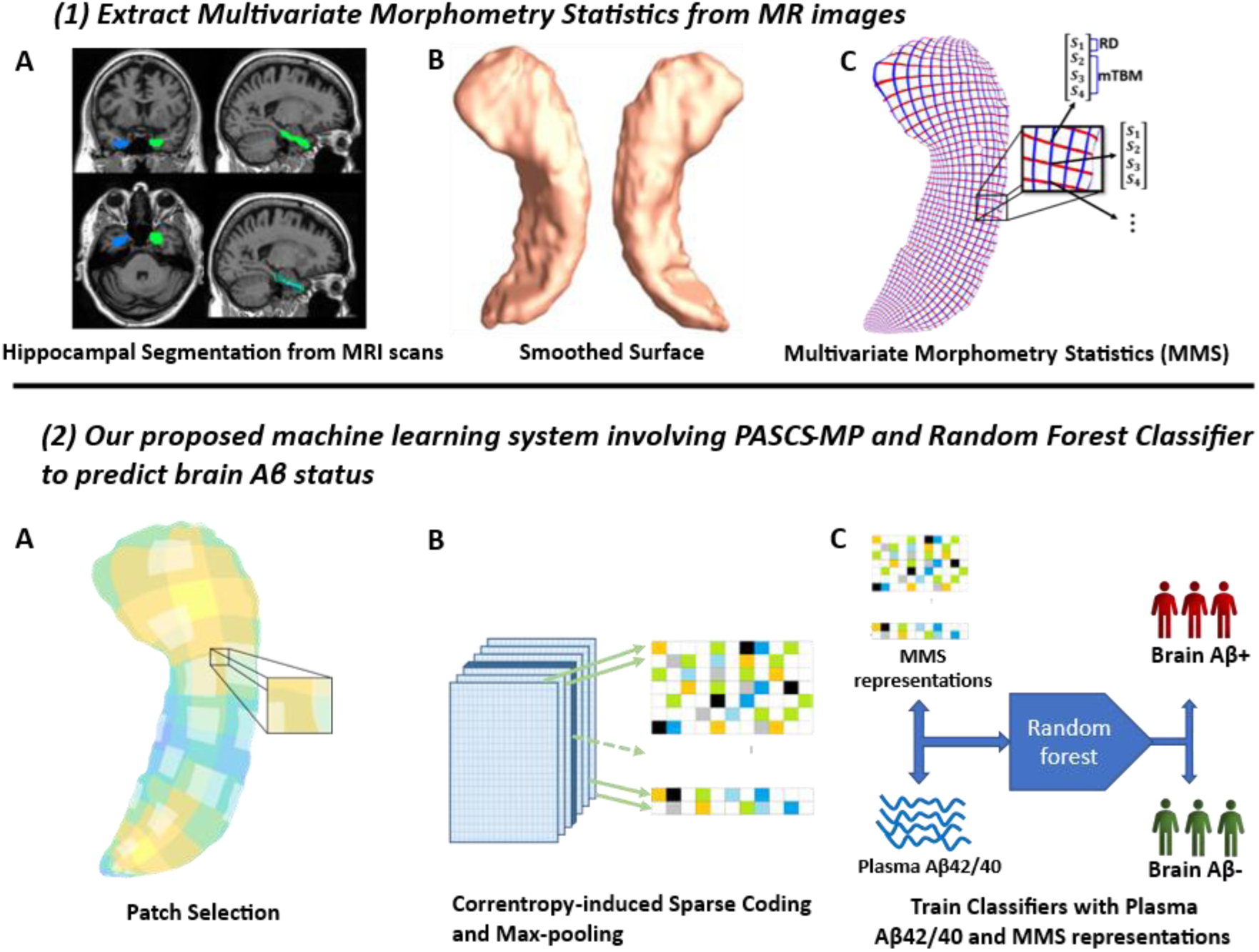
Framework for predicting brain amyloid positivity with blood-based marker Plasma Aβ42/40 and MMS-based PASCS-MP representation. Penal (1) shows MMS are extracted from MRI scans. In Penal (2), we generated representations with PASCS-MP and used them with Plasma Aβ42/40 as predictors to classify the patients with different brain amyloid positivity based on a random forest classifier.

### 2.5 Surface Feature Dimensionality Reduction

To resolve the high-dimension small sample problem, we took advantage of our previously developed Patch Analysis-based Surface Correntropy-induced Sparce Coding and Max-pooling (PASCS-MP) method to generate a representation of the subject with reasonable feature size [23]. From each hemisphere, we selected 504 patches of sizes 10*10, resulting in 1008 patches from each subject [40,50]. Then, we learned a sparse coding representation for each patch by stochastic coordinate coding algorithm, resulting in a 1008(patches)*1800(features) dimension representation for each subject as the input to max-pooling process.

We borrowed the Correntropy-induced Sparse-coding (SC) from our previous work, which improves the Stochastic Coordinate Coding (SCC) method [51]. In traditional dictionary learning approach, we learn a dictionary D and a representation Z, such that the distance between the original input X and the learned representation Z is minimized, subject to regularization on the sparsity of D:

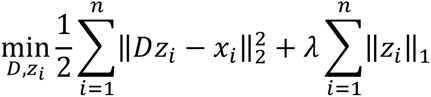

In correntropy measure, however, the similarity function can be replaced by the expectation of Gaussian kernel:

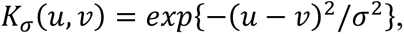

resulting in a correntropy-based loss function:

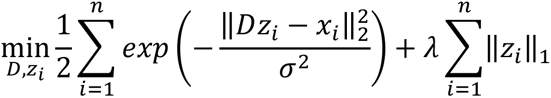

Furthermore, our recent study reformulated the first part of the equation by the half-quadratic technique and introduced auxiliary variable that can be customized according to different requirements. The final objective function becomes:

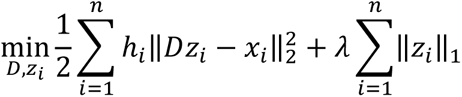

Where 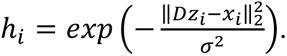

To solve the minimization problem, for each image patch, we update the dictionary D, the latent representation Z and the auxiliary variable h in turn, while keeping the other variables fixed (Algorithm 1). For details of the mathematical induction and proofs, refer to our previous study [23].

### 2.6 Max-pooling and Classification

With the computed 1008*1800-dimension representation from PASCS process, we performed max-pooling [20,52] on each feature across all the 1008 patches to keep the maximum feature values. By this process, we scaled down the dimensions of the feature vectors from 1008*1800 to 1*1800, which is a more managable size to apply traditional machine learning algorithms.

After getting the pooled image features, we separately trained three classifiers to predict the positivity of brain Aβ. First, we fit a logistic regression model of brain Aβ vs plasma Aβ42/40. The reason we used logistic regression here instead of random forest for other predictors is that plasma Aβ42/40 is a sole predictor, which is unsuitable for decision tree-based methods. Second, we trained a random forest classifier on the MMS-based features extracted by PASCS-MP pipeline described earlier. As a third step, another random forest classifier was trained using the joint features of plasma Aβ42/40 and extracted MMS. For each experiment, we performed a 10-fold cross-validation five times, and evaluated the performance based on the average accuracy, precision, recall rate and F1 score. Both logistic regression and random forest classifiers were implemented by scikit-learn package [53]. In addition, due to the imbalanced property of the BAI data, we adopt the SMOTE upsampling method to balance the positive and negative sample numbers [54].

### 2.7 Performance Evaluation Protocol

Based on our previous study, the performance of PASCS-MP pipeline relies on four key parameters: the patch size, the dimensionality of the learned representation, the regularization parameter for the l1-norm (λ) and the kernel size σ. We have previously determined that the optimal parameter values for patch size and dimensionality are 10*10 and 1800, respectively, which was verified using both ADNI and OASIS datasets [23]. In this study, we verify that the same patch size and dimensionality are still optimal for the new dataset, but the best λ and σ values for our dataset shifted to 0.4 and 2.0, respectively.

To evaluate the model performance using different feature combinations, we adopt 10-fold cross-validation by random shuffling the data samples and splitting them into ten sub-sets. We use one subset as a testing set and the other 9 subsets as training set. After classification, we generate the confusion matrix and calculate the accuracy, precision, recall rate and F1 score. We also compute the area-under-the-curve (AUC) of the receiver operating characteristics (ROC). We applied the same approach and parameter settings on both datasets.

## 3 RESULTS

### 3.1 Parameter Settings

As determined in our prior work, the best parameters for our model were sparse code dimensionality = 1800 and patch size = 10*10, regularization parameter λ = 0.22, kernel size σ = 3.6. In this work, we adopted the same dimensionality and patch size, but we sought to identify the best combination of λ and σ for our datasets, which respectively control the sparsity of the learned representation and the properties of correntropy. We examined the classification accuracy of the learned random forest classifier with 1000 nodes based solely on MMS features. The optimal parameter values were λ = 0.2, σ = 2.0 in ADNI dataset, while the best combination became λ = 0.4, σ = 2.0 in BAI dataset (Table 2). For the sake of generalization and consistency across the two datasets, we decided to use λ = 0.4, σ = 2.0 for both datasets in the subsequent experiments. Furthermore, we performed an additional search among different number of nodes in the random forest classifier and determined that the optimal number of nodes is 1000 (Table 3).

**Table 2.** Performance comparison of different combinations of λ and σ.

| Dataset | $\lambda$ | $\sigma$ | Accuracy |
| --- | --- | --- | --- |
| ADNI | **0.2** | **2** | **0.87** |
|  | 0.2 | 4 | 0.78 |
|  | 0.22 | 3.6 | 0.77 |
|  | 0.3 | 2 | 0.85 |
|  | 0.3 | 4 | 0.77 |
|  | **0.4** | **2** | **0.83** |
|  | 0.4 | 4 | 0.82 |
| BAI | 0.2 | 2 | 0.75 |
|  | 0.2 | 4 | 0.75 |
|  | 0.22 | 3.6 | 0.81 |
|  | 0.3 | 2 | 0.77 |
|  | 0.3 | 4 | 0.73 |
|  | **0.4** | **2** | **0.88** |
|  | 0.4 | 4 | 0.79 |

**Table 3.** Performance comparison of different numbers of random forest nodes in combined predictor.

| Dataset | Nodes | Accuracy | Precision | Recall | F1 score |
| --- | --- | --- | --- | --- | --- |
| ADNI | 100 | $0.77 \pm 0.01$ | $0.77 \pm 0.02$ | $0.76 \pm 0.03$ | $0.76 \pm 0.02$ |
| | 500 | $0.79 \pm 0.01$ | $0.80 \pm 0.02$ | $0.80 \pm 0.02$ | $0.80 \pm 0.01$ |
|  | **1000** | **$0.82 \pm 0.02$** | **$0.82 \pm 0.02$** | **$0.81 \pm 0.02$** | **$0.81 \pm 0.01$** |
| | 2000 | $0.81 \pm 0.01$ | $0.81 \pm 0.01$ | $0.80 \pm 0.01$ | $0.80 \pm 0.01$ |
| BAI | 100 | $0.89 \pm 0.02$ | $0.91 \pm 0.02$ | $0.86 \pm 0.02$ | $0.88 \pm 0.02$ |
| | 500 | $0.90 \pm 0.01$ | $0.92 \pm 0.01$ | $0.86 \pm 0.01$ | $0.88 \pm 0.01$ |
|  | **1000** | **$0.91 \pm 0.01$** | **$0.93 \pm 0.01$** | **$0.89 \pm 0.02$** | **$0.90 \pm 0.01$** |
| | 2000 | $0.88 \pm 0.02$ | $0.92 \pm 0.02$ | $0.85 \pm 0.03$ | $0.87 \pm 0.03$ |

### 3.2 Performance Comparison Result

#### 3.2.1 Accuracy, Precision, Recall rate, F1 score and ROC-AUC

We evaluated the performance of the three models on amyloid positivity prediction: the logistic regression model using blood Aβ42/40, the random forest model using surface MMS features, and the random forest model using the combined blood Aβ42/40 and surface MMS features. The classification accuracies using only Plasma Aβ42/40 are 0.73 in ADNI dataset and 0.81 in BAI dataset, respectively. The accuracies using only surface MMS features are 0.81 in ADNI dataset and 0.87 in BAI dataset, respectively. By integrating MMS features and plasma Aβ42/40 measurements, the model accuracy improved to 0.85 and 0.90 in ADNI and BAI datasets, respectively, outperforming either baseline models (Table 4). We also calculated the AUC scores of different feature combinations and plotted the ROC curve. As shown in Figure 2, the MMS + plasma Aβ42/40 + APOE model outperforms the others, yielding AUC scores 0.94 in ADNI dataset and 0.98 in BAI dataset respectively. We performed Delong test to evaluate the improvement of AUC by incorporating imaging features, APOE genotype, age and sex into our model. As a result, the integrated model achieved a significant improvement compared with plasma Aβ42/40 predictor (p-value = 0.0024 in ADNI and 0.0021 in BAI), but the improvement was not statistically significant over the image-only model (p-value = 0.59 in ADNI and 0.99 in BAI). In addition, the models with or without covariates yield similar performance (p-value = 0.28-0.95 in both datasets). Moreover, we evaluated the model with highest classification accuracy, i.e., MMS + Aβ42/40 + APOE mode, on different diagnosis groups in ADNI dataset. The model performance was stable across diagnosis groups (Accuracy = 0.90 in CN group and 0.89 in MCI group).

**Figure 2.**
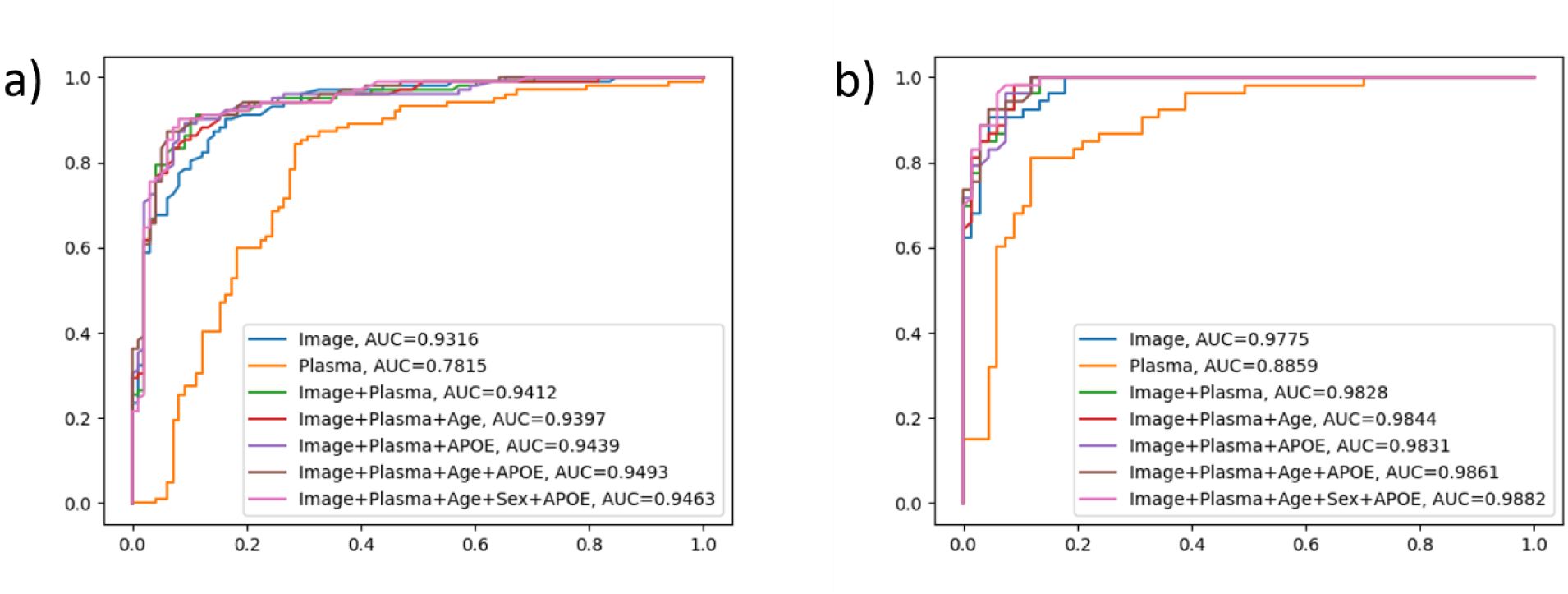
ROC curves of classification models with different feature combinations. a) ROC curves of ADNI dataset. b) ROC curves of BAI dataset. Incorporated models with image MMS and plasma Aβ42/40 features outperforms the individual models using only image MMS features or plasma Aβ42/40 as predictors. Adding covariates slightly improves the AUC score.

**Table 4.** Performance comparison of different predictors.

| Dataset | Feature | Accuracy | Precision | Recall | F1 score |
| --- | --- | --- | --- | --- | --- |
| ADNI | Plasma Aβ42/40 (Logistic) | $0.73 \pm 0.01$ | $0.73 \pm 0.01$ | $0.73 \pm 0.01$ | $0.73 \pm 0.01$ |
| | MMS (RF) | $0.81 \pm 0.01$ | $0.81 \pm 0.01$ | $0.80 \pm 0.01$ | $0.81 \pm 0.01$ |
| | MMS + Aβ42/40 (RF) | $0.85 \pm 0.01$ | $0.84 \pm 0.01$ | $0.84 \pm 0.01$ | $0.84 \pm 0.01$ |
| | MMS + Aβ42/40 + Age (RF) | $0.84 \pm 0.01$ | $0.84 \pm 0.01$ | $0.83 \pm 0.02$ | $0.83 \pm 0.01$ |
|  | **MMS + Aβ42/40 + APOE (RF)** | **$0.86 \pm 0.01$** | **$0.85 \pm 0.02$** | **$0.84 \pm 0.02$** | **$0.85 \pm 0.01$** |
| | MMS + Aβ42/40 + Age + APOE (RF) | $0.83 \pm 0.01$ | $0.85 \pm 0.02$ | $0.85 \pm 0.02$ | $0.85 \pm 0.02$ |
| | MMS + Aβ42/40 + Age + Sex + APOE (RF) | $0.85 \pm 0.01$ | $0.85 \pm 0.01$ | $0.85 \pm 0.01$ | $0.85 \pm 0.01$ |
| BAI | Plasma Aβ42/40 (Logistic) | $0.81 \pm 0.01$ | $0.80 \pm 0.01$ | $0.81 \pm 0.01$ | $0.80 \pm 0.01$ |
| | MMS (RF) | $0.87 \pm 0.01$ | $0.90 \pm 0.01$ | $0.83 \pm 0.02$ | $0.85 \pm 0.02$ |
| | MMS + Aβ42/40 (RF) | $0.90 \pm 0.02$ | $0.92 \pm 0.01$ | $0.87 \pm 0.02$ | $0.89 \pm 0.02$ |
| | MMS + Aβ42/40 + Age (RF) | $0.90 \pm 0.01$ | $0.92 \pm 0.01$ | $0.87 \pm 0.01$ | $0.89 \pm 0.01$ |
|  | **MMS + Aβ42/40 + APOE (RF)** | **$0.92 \pm 0.01$** | **$0.91 \pm 0.02$** | **$0.86 \pm 0.03$** | **$0.87 \pm 0.02$** |
| | MMS + Aβ42/40 + Age + APOE (RF) | $0.88 \pm 0.01$ | $0.92 \pm 0.01$ | $0.87 \pm 0.02$ | $0.88 \pm 0.02$ |
| | MMS + Aβ42/40 + Age + Sex + APOE (RF) | $0.89 \pm 0.01$ | $0.92 \pm 0.01$ | $0.87 \pm 0.01$ | $0.89 \pm 0.02$ |

We also evaluated the influence of individual demographic features, including age and sex, by incorporating them into the logistic regression model and testing for significance. As a result, both age and sex showed significance (p-value < 0.05) in the univariate models in ADNI, while sex was not significant in BAI dataset (Table 5). Therefore, we evaluated age and sex as covariates in our combined models. A complete list of all combinations and corresponding performance are listed in Table 4. Overall, we observed that incorporating age slightly reduced the standard deviation of the performance statistics, but the classification accuracies were not substantially influenced.

**Table 5.** Univariate logistic regression of potential covariates. Significance Code: 0: ‘***’; 0.001: ‘**’; 0.01: ‘*’; 0.05: ‘.’

|  | p-value (ADNI) | p-value (BAI) |
| --- | --- | --- |
| Age | 0.004 ** | 0.002 ** |
| Sex | 0.036 * | 0.168 |

## 4 DISCUSSION

In this study, we assessed a novel approach for detecting brain amyloid-PET positivity using an integrated multimodal model that combines an MR image feature and a plasma Aβ42/40 BBBM. To the best of our knowledge, this is the first study that combines this type of advanced morphometric brain imaging information and amyloid BBBMs to assess amyloid-PET positivity. Our previously developed PASCS-MP method for geometric feature extraction successfully generated features that achieved accuracy of 0.81 ± 0.01 in ADNI dataset and 0.87 ± 0.01 in BAI dataset with a random forest classifier based on MR data alone. With the integration of plasma Aβ42/40 measures, we further improved the accuracy to 0.86 ± 0.01 and 0.92 ± 0.01 in the two datasets respectively, surpassing both baseline unimodal models that used only a single AD-related feature. This result verified our hypothesis that the MMI classification model that integrated MRI image features and plasma biomarkers can achieve superior performance compared to each individual unimodal model. The slight difference in accuracy for plasma Aβ42/40 assay alone observed in this study compared to the original studies [29,31] is likely due to the differences in imaging analysis methods and the determination of PET based amyloid positivity.

In our analysis, although univariate logistic regression showed a significant *p*-value for age, adding age or sex as a lone covariate to the IMM model did not improve accuracy. However, when age and sex data were both included AUCs improved AUC in both datasets suggesting a potential interaction effect that could enhance model robustness. Adding APOE genotype information to the MMI model also improved performance, suggesting that APOE genotype can provide additional predictive power beyond our structural MR feature and plasma Aβ42/40. Further investigations with larger and more diverse datasets would provide additional insights into the potnetial utility of including these covariates in predicting brain amyloid-PET positivity.

A study limitation involved the moderate sample size (198 for the ADNI dataset and 111 for the BAI dataset) which has the potential to introduce dataset-specific parameters and result in overfitted models. To mitigate this limitation, we made an effort to avoid selecting separate λ and σ model parameter values for each dataset. Instead, we picked a shared set of λ and σ that achieved high prediction accuracy across the two datasets in our preliminary parameter search (Table 2). Even so, the parameter values differ from the determined optimal values in our previous study using larger datasets [23]. With more data available in the future, we expect a more stable and robust parameter setting that may be more generalizable across heterogeneous datasets. Another limitation of the study is that we used a specific plasma Aβ-related assay while there are now a range of other BBBMs, including plasma tau phosphorylated at different sites (pTau181, pTau217, pTau231), that show promise for predicting AD-cerebral amyloidosis [28,55,56]. For example, for AD progression prediction, Tau217 and pTau181, combined with memory, executive function and APOE, achieved high prediction accuracy in BioFINDER (AUC = 0.91) and ADNI (AUC = 0.90) cohorts, respectively [57]. Further studies are warranted to examine the combined performance of our MR based approach together with other BBBMs in the detection of amyloid status across the AD continuum.

In summary, these results suggest the potential of advanced structural MRI features providing similar accuracy to plasma Aβ42/40 in classifying amyloid-PET positivity, and support the promise of integrating such advanced structural MR features with blood-based biomarkers along with demographic information and APOE4 status can further improve amyloid-PET predictive accuracy. These multimodal and multiplex integrated biomarker approaches hold great promise and merit further investigation to aid improvements in developing and utilizing more efficient and accurate AD biomarkers for clinical and research use, and to advance therapeutic and preventative strategies.

## 5 ACKNOWLEDGMENTS

The ADNI Data used in the preparation of this article was obtained from the Alzheimer’s Disease Neuroimaging Initiative (ADNI) database (adni.loni.usc.edu). The ADNI was launched in 2003 as a public-private partnership led by Principal Investigator Michael W. Weiner, MD. The primary goal of ADNI has been to test whether serial MRI, PET, other biological markers, and clinical and neuropsychological assessments can be combined to measure the progression of MCI and early AD. For up-to-date information, see www.adni-info.org.

Data collection and sharing for this project was funded by the Alzheimer’s Disease Neuroimaging Initiative (ADNI) (National Institutes of Health Grant U01 AG024904) and DoD ADNI (Department of Defense award number W81XWH-12-2-0012). ADNI is funded by the National Institute on Aging, the National Institute of Biomedical Imaging and Bioengineering, and through generous contributions from the following: Alzheimer’s Association; Alzheimer’s Drug Discovery Foundation; BioClinica, Inc.; Biogen Idec Inc.; Bristol-Myers Squibb Company; Eisai Inc.; Elan Pharmaceuticals, Inc.; Eli Lilly and Company; F. Hoffmann-La Roche Ltd and its affiliated company Genentech, Inc.; GE Healthcare; Innogenetics, N.V.; IXICO Ltd.; Janssen Alzheimer Immunotherapy Research & Development, LLC.; Johnson & Johnson Pharmaceutical Research & Development LLC.; Medpace, Inc.; Merck & Co., Inc.; Meso Scale Diagnostics, LLC.; NeuroRx Research; Novartis Pharmaceuticals Corporation; Pfizer Inc.; Piramal Imaging; Servier; Synarc Inc.; and Takeda Pharmaceutical Company. The Canadian Institutes of Health Research is providing funds to support ADNI clinical sites in Canada. Private sector contributions are facilitated by the Foundation for the National Institutes of Health (www.fnih.org). The grantee organization is the Northern California Institute for Research and Education, and the study is coordinated by the Alzheimer’s Disease Cooperative Study at the University of California, San Diego. ADNI data are disseminated by the Laboratory for Neuro Imaging at the University of Southern California.

## 6 Code Availability

Source code of this paper is available on github: https://github.com/ychen855/PASCS-MP.

## 7 Data Availability

Data used in this study are available from the corresponding author, Yalin Wang, upon reasonable request.

## 8 APPENDIX

### Algorithm 1. Patch analysis-based surface correntropy-induced sparse-coding

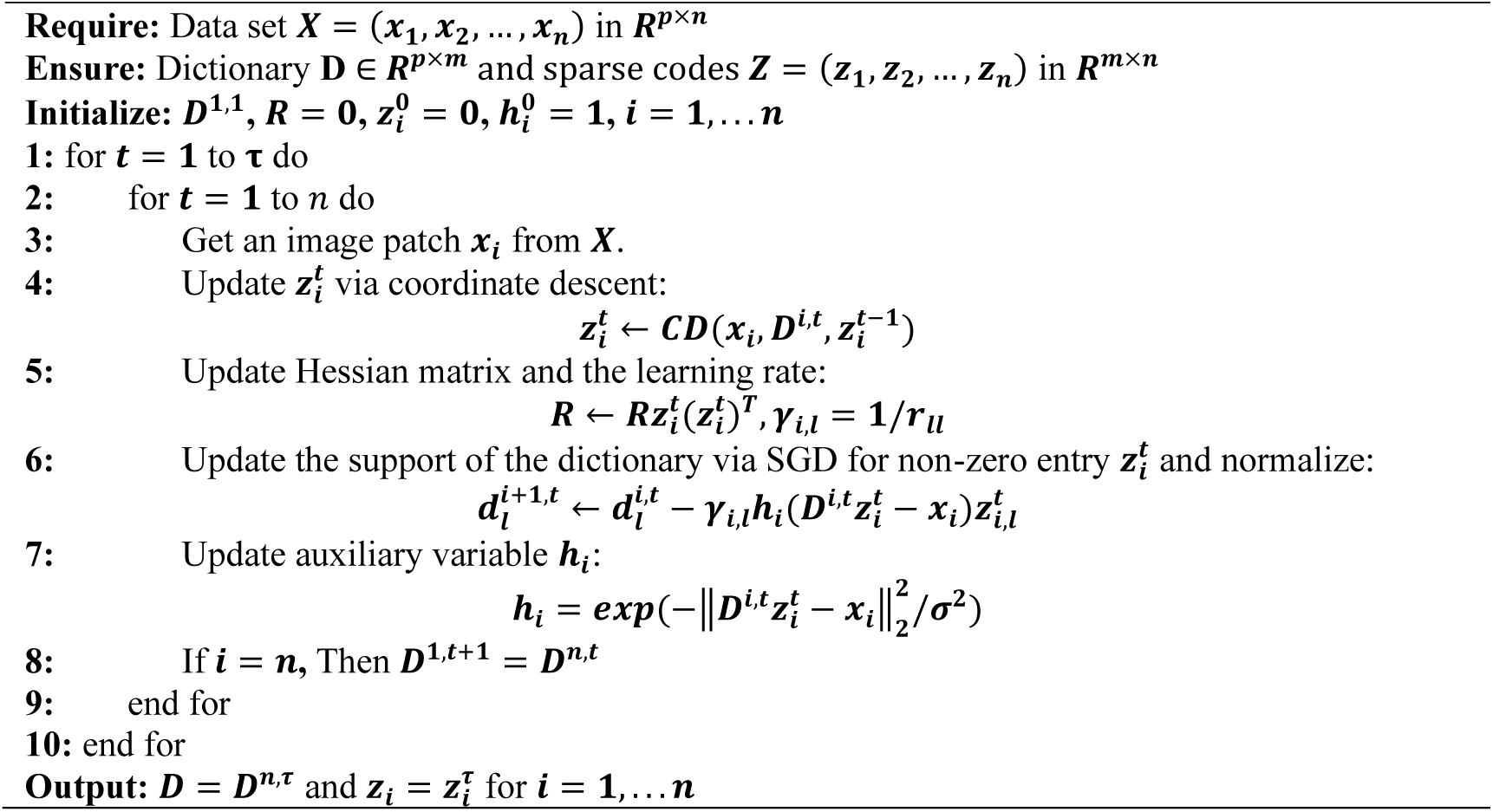

